# PPIDomainMiner : Inferring domain-domain interactions from multiple sources of protein-protein interactions

**DOI:** 10.1101/2021.03.03.433732

**Authors:** Seyed Ziaeddine Alborzi, Amina Ahmed Nacer, Hiba Najjar, David W Ritchie, Marie-Dominique Devignes

**Author notes:** EcoMundo, Issy-les-Moulineaux, F-92130, France (Seyed Ziaeddine Alborzi). Deceased (David W Ritchie).

## Abstract

Many biological processes are mediated by protein-protein interactions (PPIs). Because protein domains are the building blocks of proteins, PPIs likely rely on domain-domain interactions (DDIs). Several attempts exist to infer DDIs from PPI networks but the produced datasets are heterogeneous and sometimes not accessible, while the PPI interactome data keeps growing.

We describe a new computational approach called “PPIDM” (Protein-Protein Interactions Domain Miner) for inferring DDIs using multiple sources of PPIs. The approach is an extension of our previously described “CODAC” (Computational Discovery of Direct Associations using Common neighbors) method for inferring new edges in a tripartite graph. The PPIDM method has been applied to seven widely used PPI resources, using as “Gold-Standard” a set of DDIs extracted from 3D structural databases. Overall, PPIDM has produced a dataset of 84, 552 non-redundant DDIs. Statistical significance (p-value) is calculated for each source of PPI and used to classify the PPIDM DDIs in Gold (9,175 DDIs), Silver (24, 934 DDIs) and Bronze (50, 443 DDIs) categories. Dataset comparison reveals that PPIDM has inferred from the 2017 releases of PPI sources about 46% of the DDIs present in the 2020 release of the 3did database, not counting the DDIs present in the Gold-Standard. The PPIDM dataset contains 10, 229 DDIs that are consistent with more than 13, 300 PPIs extracted from the IMEx database, and nearly 23,300 DDIs (27.5%) that are consistent with more than 214,000 human PPIs extracted from the STRING database. Examples of newly inferred DDIs covering more than 10 PPIs in the IMEx database are provided.

Further exploitation of the PPIDM DDI reservoir includes the inventory of possible partners of a protein of interest and characterization of protein interactions at the domain level in combination with other methods. The result is publicly available at http://ppidm.loria.fr/.

**Author summary:** We revisit at a large scale the question of inferring DDIs from PPIs. Compared to previous studies, we take a unified approach accross multiple sources of PPIs. This approach is a method for inferring new edges in a tripartite graph setting and can be compared to link prediction approaches in knowledge graphs. Aggregation of several sources is performed using an optimized weighted average of the individual scores calculated in each source. A huge dataset of over 84K DDIs is produced which far exceeds the previous datasets. We show that a significant portion of the PPIDM dataset covers a large number of PPIs from curated (IMEx) or non curated (STRING) databases. Such a reservoir of DDIs deserves further exploration and can be combined with high-throughput methods such as cross-linking mass spectrometry to identify plausible protein partners of proteins of interest.

## Introduction

Many biological processes from metabolic pathways to cellular signaling are mediated by protein-protein interactions (PPIs). However, the experimental determination and analysis of such interactions are often difficult and time-consuming. The developments in high-throughput gene sequencing techniques have created a huge gap between the ever increasing number of known protein sequences and the knowledge of their function and of their biological interactions. There is therefore much interest in developing computational approaches to predict PPIs. Because protein domains are the building blocks of proteins, PPIs mainly rely on given combinations of domain-domain interactions (DDIs). Predicting and modelling PPIs should therefore benefit from a systematic inventory of all possible DDIs.

In fact, the interest raised by DDI study is justified by simple combinatorial reasoning. As most proteins are composed of a limited subset of domains, the study of all possible interactions between the hundreds of millions of proteins available in the sequence databases can be advantageously reduced by the study of the possible interactions between the tens of thousands of domains existing in current domain classifications such as Pfam (17,929 families in the 2018 release 32; [1]) or CDD (Conserved Domain Database, 52,910 conserved domain models in the v3.17 version; [2]. It is therefore of utmost importance to enumerate all plausible DDIs which could explain all the PPIs described so far and which could be used to predict interactions which have not yet been described.

Today, well established DDI resources are based on experimental evidence such as 3D structures extracted from the Protein Data Bank (PDB). This is the case of 3did (updated every year; [3–5]), KBDOCK (last release in 2016; [6]), iPfam (last release in 2013; [7]) and INSTRUCT (last release in 2013; [8]). However the DDI content of these databases does not account for all PPIs described so far. For example, it has been estimated in 2015 that DDIs extracted from 3did only cover around 20% of PPIs from the STRING database [9].

For this reason, many computational methods have been proposed for inferring DDIs from the domain composition of proteins involved in PPIs. The pioneer work by Sprinzak and Margalit in 2001 used a simple correlation measure of domain occurrence in PPIs [10]. It was rapidly extended and improved in quite diverse manners.

A first group of methods uses probabilistic or statistical approaches including measures of the interaction probability between two domains [11,12] or involving expectation maximization algorithm to maximize a certain likelihood function over a given interactome [13–15], enrichment calculation [16], sometimes with the help of functional (Gene Ontology) annotations [17,18] or with complex network neighborhood coefficients [9].

Other methods exploit the properties of DDIs related to sequence and gene evolution, such as the concepts of domain fusion [19], phylogenetic profiling [20], relative co-evolution of domain pairs [21,22], correlated mutations [23], or combination of these properties [24].

Another group of methods aims to explain a given set of PPIs with a minimum set of DDIs using various optimization methods such as Integer Linear Programming for parsimonious explanation [25–27], genetic algorithm [28], or parameter-dependent selection [29].

Finally, other authors use PPI and non PPI datasets to develop machine learning approaches such as random forest framework [30,31], discriminant feature selection methods [32], or formal concept analysis [33].

The various datasets of inferred DDIs generated by all these methods do not overlap perfectly well, mainly because they rely on various non overlapping protein interactomes such as DIP (Database of Interacting Proteins; [34]), IntAct [35], HPRD (Human Protein Reference Database; [36]) or STRING [37]. Thus, several attempts have tried to merge a certain number of the produced DDI datasets and compute a score for each DDI depending on its presence in the composing resources.

The most famous integrated resource is certainly the DOMINE database, created in 2008 and updated in 2011 [38,39]. In its last available release (2011), DOMINE has merged not less than 12 sources of inferred DDIs with two sources of DDIs derived from 3D structures, namely iPfam (2007 release) and 3did (2010 release), leading to a dataset of 26,219 DDIs involving 5,140 distinct Pfam domains.

The UniDomInt [40] and IDDI (Integrated DDI; [41]) resources are other initiatives aimed at merging well-established sets of structural DDIs with sets of inferred DDIs from various methods (9 and 20 sets, respectively). The DIMA3.0 database (Domain Interaction MAp; [23]) integrates more than 5,800 structurally known DDIs from iPfam and 3did databases and 46,900 DDIs computed with four of the methods listed above.

The fact that different methods infer the same DDIs can be considered as an argument for DDI reliability. Indeed, in the DOMINE resource, an index quantifying the percentage of overlap (POI) between the twelve sources has been defined and each DDI has received a score calculated as the sum of the POIs of all the sources inferring this DDI. Depending on their score and on their participation in the same biological process (according to the Lee’s dataset; [17]), DDIs are classified as high, medium or low confidence. Resources such as UniDomInt and IDDI use slightly different methods to assess consistency between the DDIs datasets [40, 41].

The extreme heterogeneity of methods for inferring DDIs and the absence of regular updates for most of the generated datasets have lead to the absence of any available large-scale updated repository of scored DDIs, reflecting the multiple PPI resources available today. Such a repository could be advantageous in studying PPIs at the domain level.

Therefore, we were encouraged to develop a generic new method capable of exploiting multiple PPI datasets for the automatic inference of DDIs. The method is called PPIDM (for Protein-Protein Interaction Domain Miner) and is inspired both by the statistical approaches described above and by the CODAC method, that we previously designed to infer protein domain annotations [42]. CODAC uses a tri-partite graph setting and a Gold-Standard of existing associations to infer all possible associations between two sets of objects, assuming the objects share a similar neighborhood in a third set of objects. In PPIDM, the two sets of objects are sets of domains and the third set is a set of PPIs.

In this paper, we first describe the PPIDM algorithm and its application to seven datasets of PPIs using a Gold-Standard of 7, 254 DDIs derived from 3did and KBDOCK. We then compare the inferred DDIs (84,552 pairs of Pfam domains) with inferred DDIs in DOMINE, revealing a large percentage of newly inferred DDIs in PPIDM. We also analyse the overlap with the 2017 to 2020 releases of 3did, revealing that the DDIs inferred by PPIDM overlap with 3, 239 (46%) DDIs present in the 3did 2020 release (not counting the Gold-Standard), including 800 newly observed DDIs that were not known in 3did when PPIDM was generated. We also study the coverage of PPIs from curated (IMEx) or non curated (STRING) databases by PPIDM DDIs and we compare with DOMINE to the advantage of PPIDM. Finally, we describe the biological plausability of a subset of newly inferred DDIs from PPIDM.

## Materials and methods

### CODAC-inspired PPIDM approach

The CODAC-inspired PPIDM approach is based on a tripartite graph setting [42]. In graph theory, a *k*-partite graph is a graph whose vertices can be partitioned into *k* disjoint subsets, such that in each subset no two vertices are connected. If *k* = 2, the graph is called a bipartite graph (or bigraph), and if *k* = 3 it is called a tripartite graph. The tripartite graph setting is designed here to solve problems of bipartite graph enrichment also known as edge prediction or inference. The main intuition is to calculate new weighted edges between two sets of items when sparse known edges already exist between the two and when both sets display dense connections with a common third set of items. Let 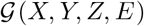 be a tripartite graph where *X*, *Y* and *Z* are 3 sets of items and *E* is the set of all edges connecting *X*, *Y* and *Z* in the input configuration. Let us consider the 3 bipartite subgraphs of 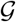, denoted as 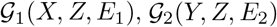, and 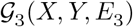. We now assume that the set of edges *E*_3_ is incomplete, and that the aim is to compute new edges between items of *X* and items of *Y* in order to generate 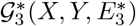 which together with 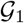 and 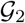 will make the final tripartite graph, 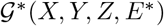, where *E*\* denotes an enriched set of edges. New edges may be inferred by exploiting the existing edge distributions in 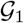 and 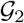. The hypothesis here is that if items *x_i_* of *X* and *y_j_* of *Y* share the same (or almost the same) set of neighbors {*z_k_*} in *Z*, then it may be inferred, with a certain confidence to be estimated, that an edge exists between *x_i_* and *y_j_*.

In the PPIDM approach, we want to infer new DDIs between domains that are present in the proteins participating in PPIs. Thus, we instantiate the tripartite graph settings with *Z*, a set of PPIs, and *X* and *Y*, two sets of Pfam domains that compose the proteins from *Z*, as shown in Fig 1. More precisely, we distinguish the left and right components of the pair of proteins forming a PPI and ordered according to the alphanumerical order of their identifiers (Ids). The first bipartite graph 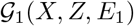 represents the relations between domains of *X* and the left proteins of PPIs in *Z*, while the second bipartite graph 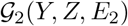 represents the relations between domains of *Y* and the right proteins of PPIs in *Z*. In Fig 1, the *X* and *Y* sets are named *D_L_* and *D_R_*, respectively. By construction, *D_L_* and *D_R_* are overlapping but distinct sets of domains encompassing the domains present in the left and right proteins of PPIs, respectively. The third bipartite graph 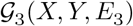 is initialized with a set of edges representing DDIs observed in structural databases. Our Gold-Standard set of edges *E*_3_ pertains from the intersection of the KBDOCK and 3did databases.

**Fig 1.**
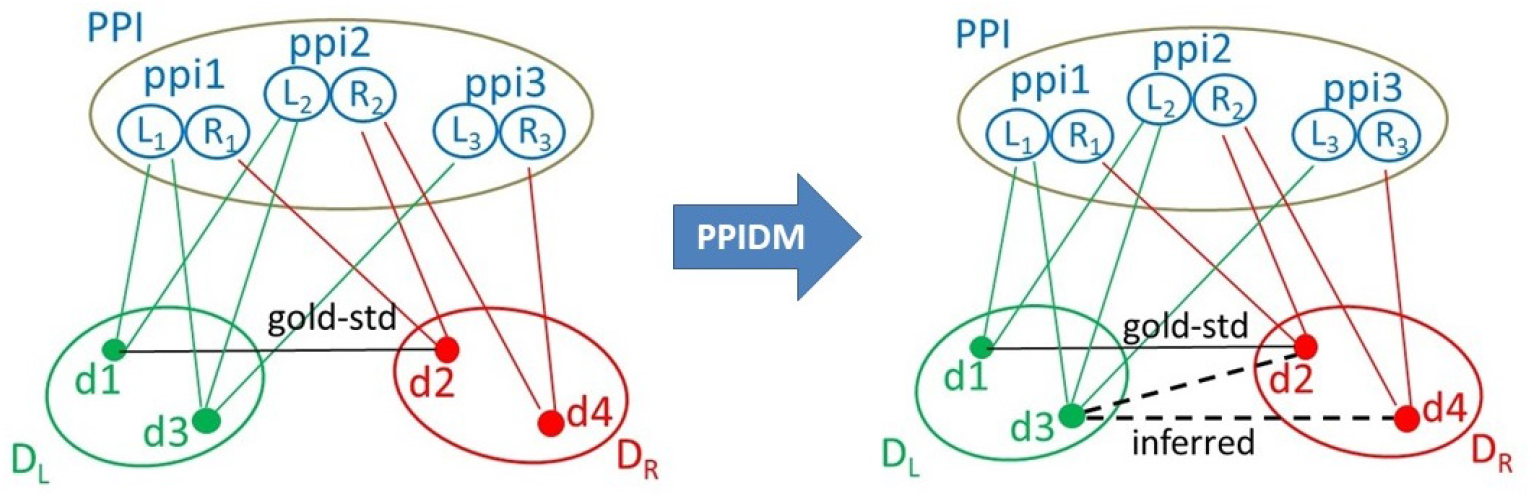
Schematic illustration of edge inference by PPIDM in a tripartite graph setting 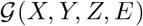. *Z* is here PPI, a set of PPIs, *X* and *Y* are *D_L_* and *D_R_*, two sets of Pfam domains. Each item in *PPI* is an ordered pair of proteins *ppi_i_* = (*L_i_, R_i_*) with *Id*(*L_i_*) ≤ *Id*(*R_i_*). Domains in *D_L_* and *D_R_* are connected to their common neighbor item *ppi_i_* in PPI through *L_i_* and *R_i_* respectively. The (d1, d2) edge comes from the Gold-Standard dataset of DDIs. With PPIDM, new edges are inferred between domains of *D_L_* and domains of *D_R_* if their adjacency vectors in *PPI* are similar. Here, the (d3, d2) edge is inferred because d3 and d2 are found in ppi1 and ppi2, and (d3, d4) is inferred because d3 and d4 are found in ppi2 and ppi3. However, the score of (d3, d2) will be lower than the score of (d1, d2) because d3 has one neighbor that does not contain d2 (namely ppi3).

The output of the PPIDM algorithm is 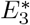, which will contain an enriched set of DDIs weighted by their neighborhood similarity score. The adjacency matrices *M_X_* of *X* in *Z* and *M_Y_* of *Y* in *Z* are built and cosine similarities are computed between each row of *M_X_* and each row of *M_Y_*, representing the neighborhood similarity score of the corresponding two domains in the PPI dataset *Z*.

At this stage of the work a p-value can be calculated to estimate the significance of inferring a certain edge (*x, y*) given *Z*. Indeed, it is reasonable to suppose that an edge (*x, y*) could get a high score at random if the *x* and *y* items are each highly connected to many items in *Z*. Here, we assume that the probability of finding an edge (*x, y*) by random chance is given by an hypergeometric distribution of the number of common neighbors (*x, z*) and (*y, z*). Letting *N_z_* denote the total number of items in *Z*, *N_x_* the number of neighbors of *x* in *Z*, and *N_y_* the number of neighbors of *y* in *Z*, the probability that the (*x, y*) pair gets a number of common neighbors *K* greater than or equal to the observed *K_x,y_* is given by the hypergeometric probability equation (1):

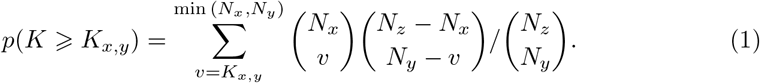

Because the p-value test is calculated for a large number of (*x, y*) edges in 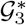, we apply a Bonferroni correction to take into account the so-called family-wise error rate [43]. Therefore, letting 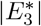 denote the total number of edges tested, we consider any p-value less than 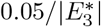 as denoting a statistically significant edge.

In order to determine an edge similarity threshold, we need to define a learning set of positive and negative examples of DDIs. Here, we take all the Gold-Standard DDIs as positive examples (*Pos*). In practice, we extract a total of 8,581 and 8,670 DDIs from 3did and KBDOCK, respectively. We then obtain 7,254 common DDIs in which each domain is represented by a Pfam identifier. These DDIs are scored among others in the general cosine similarity matrix associated with the tripartite graph. To create negative examples, we shuffle the edges corresponding to the *Pos* DDIs in 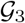 in order to rearrange in a random way all the Gold-Standard DDIs, while keeping the node degrees of each *x_i_* and each *y_j_* conserved, and not allowing to return to the original edges. We then select at random 7,254 shuffled negative DDIs and look for their score in the general cosine similarity matrix. We could get a score for only 5,145 of them (named thereafter the *Neg* DDIs) but imbalanced data is not a problem here since we used *F*_1_ – *score* for selecting the threshold.

Taken together, the *Pos* and *Neg* DDIs constitute our learning set for threshold determination. We randomly split the learning set into two groups with equal proportions of positive and negative examples to give the “Training” and “Test” subsets representing respectively two- and one-third of the learning set. We then apply 5-fold cross-validation to the Training subset to determine the optimal score threshold leading to maximal F-measure. For each fold, we rank the scores of 4/5 folds and label them “positive” or “negative” according to a score threshold that is varied in three phases with increasing resolution : firstly from 0.00 to 1.00 in steps of 0.01, then from 0.00 to 0.04 in steps of 0.001, and finally from 0.01 to 0.02 in steps of 0.0001. This allows us to determine the numbers of true positive (*TP*), false positive (*FP*), true negative (*TN*), and false negative (*FN*) predictions for each threshold value. We then calculate the recall, *R* = *TP*/(*TP* + *FN*), precision, *P* = *TP*/(*TP* + *FP*), and *F*_1_ – *score*, *F*_1_ = 2*RP*/(*P* + *R*). The similarity threshold *T* that gives the best *F*_1_ – *score* in the 4/5 folds is verified using the fifth fold, the procedure is repeated for the five folds taken as test fold. The mean threshold *T_m_* is then used to calculate the resulting *F*_1_ – *score* on the entire Training set. The robustness of this threshold is ultimately verified by calculating the *F*_1_ – *score* on the Test set (not used during threshold optimization). If satisfying, our approach is validated and the mean threshold is retained to calculate a filtered cosine similarity matrix, *C*\*, according to 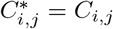 if *C_i,j_* > *T_m_*, and 0 otherwise.

### Combining Multiple Datasets

One original feature of the PPIDM approach is to be designed for handling multiple data sources of PPIs (*Z* in our tripartite graph model). When more than one PPI datasource is used, a tripartite graph is built for each data source *d* and processed separately to calculate its respective cosine similarity matrix *C^d^*. All matrices are then combined in a unique consensus matrix merging all domain subsets considered in all the graphs. Whenever there is no data for a given pair (*x, y*) in an input graph 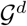, the corresponding score 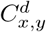 is set to zero. The cosine similarity scores are then combined as a non-zero weighted average (over all non-zero scores) to give a consensus similarity matrix, *CS*.

Weight determination is performed using Receiver-Operator-Characteristic (ROC) analysis as for information retrieval system when one wishes to retrieve positive documents as first ranked, *i.e*. with the best scores [44]. Here the positive examples are taken from the Gold-Standard set of positive DDIs and the background will be formed by all the scored DDIs in the consensus similarity matrix. One advantage of ROC-based approaches is that they are rather insensitive to the particular number of positive and negative instances used [46]. Thus, in order to find the best values for the weights *w_d_* relative to each data source used, each weight is varied from 0.01 to 1 in steps of 0.01. For each combination of weights, a ROC performance curve is calculated using the complete ranked list of consensus scores and our Gold-Standard set of positive examples. The combination of weights that gives the largest area under the curve (AUC) is selected and used to calculate the best consensus similarity matrix *CS*.

Algorithm 1 summarizes all this procedure that allows PPIDM to handle multiple data sources of PPIs to infer DDIs.

We finally classify our DDIs into *Gold, Silver*, and *Bronze* categories using the p-values calculated for each DDI in each input databases. *Gold* DDIs are those with a non-zero score in at least half of the data sources and that display a significant p-value in all these data sources, *Silver* DDIs are those with a non-zero score in less than half of the data sources while displaying a significant p-value in all these data sources, and *Bronze* are the remaining DDIs.

### Application to PPI sources

In this study, seven PPI sources have been used: IntAct and MINT [47], DIP [34], HPRD [36], and BioGRID [48,49] are manually curated databases of physical interaction between proteins; the very extensive STRING database [37] contains both physical and predicted interactions between proteins; the SIFTS database [50, 51] allows to access to PPIs present as complexes between PDB chains in the PDB. Thus, these seven PPI databases together provide a comprehensive combination of all available protein interactions and are appropriate for our PPIDM approach. Note that we retrieve all available interactions from these databases and we do not discriminate between stable and transient interactions.

#### Algorithm 1 Calculating a Consensus Similarity Matrix from Multiple Sources of Common Neighbors

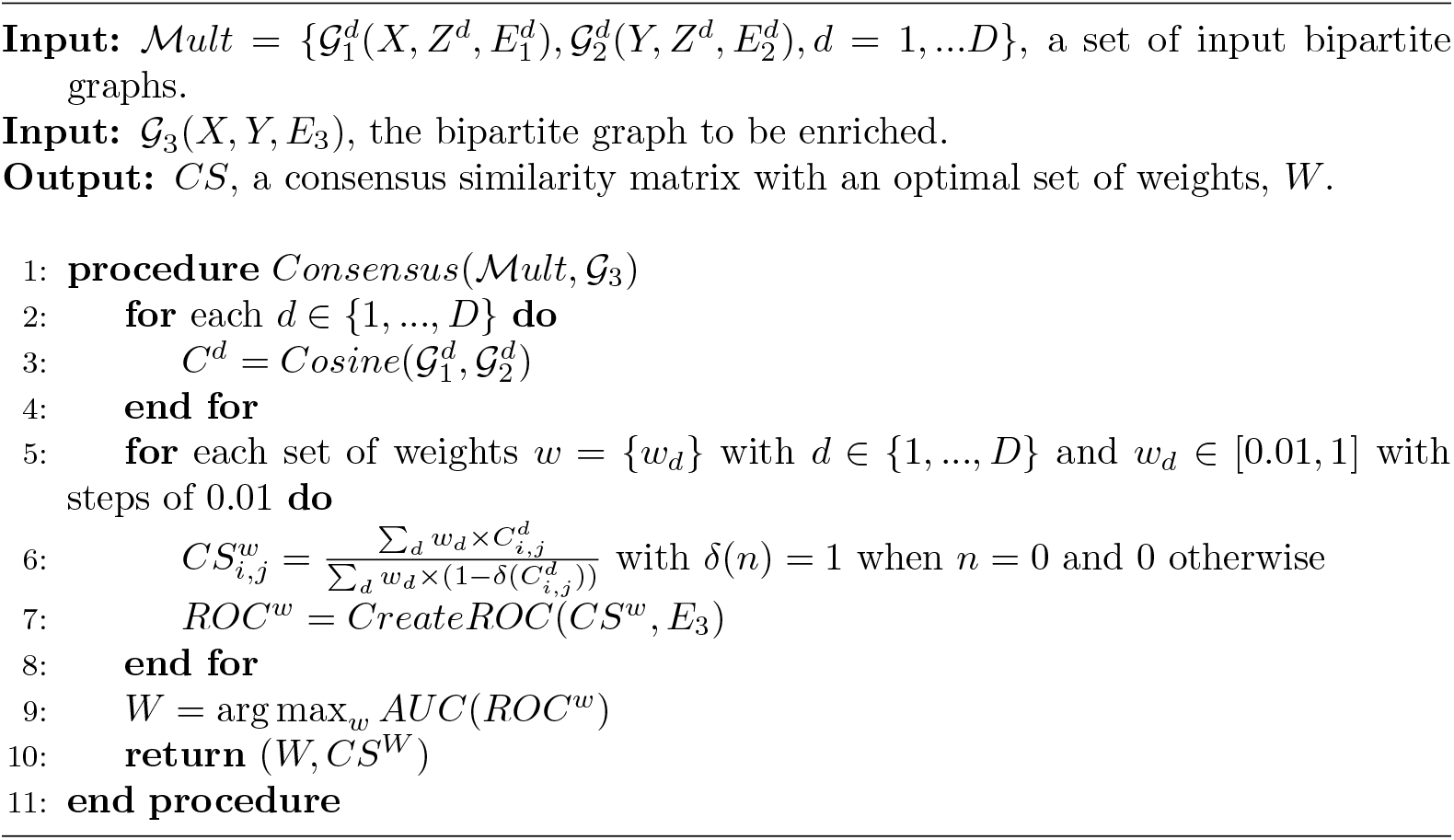

Flat data files of IntAct, DIP, MINT, HPRD, BioGRID, STRING, SIFTS, KBDOCK, 3did and UniProt (all releases as of February 2017), were downloaded and parsed using in-house Python scripts. The four sources IntAct, DIP, MINT, HPRD and SIFTS contained PPIs expressed as ordered pairs of UniProt sequence accession numbers (UniProtIds). In BioGRID and STRING, proteins are designated with an identifier system specific to each database. These identifiers were mapped to UniProtIds to produce the BioGRID and STRING datasets of PPIs. In the SIFTS database, associations between PDB chains were extracted and PPIs displaying a high possibility of interaction according to [52] were stored. Then, PDB chains possessing a representative UniProtId were replaced by this UniProtId.

The numbers of PPIs obtained from the input resources are shown in Table 1. A very large number of protein interactions is drawn from the STRING database, while the SIFTS database only provides a small collection of observed PPIs. We then categorized the STRING PPIs according to experimental and non-experimental (Text mining, Neighborhood, Fusion-fission events, Occurrence, and Coexpression) labels and stored them as STRING-exp and STRING-rest datasets, respectively.

**Table 1.** Number of interactions, distinct sequences and Pfam domains obtained from the IntAct, MINT, DIP, HPRD, BioGRID, STRING, and SIFTS (all releases as of February 2017

|  | Number of |  |  |
| --- | --- | --- | --- |
|  | PPIs | Protein Sequences | Associated Pfams |
| IntAct | 411,624 | 76,747 | 9,898 |
| MINT | 67,191 | 24,735 | 6,438 |
| DIP | 53,585 | 20,489 | 6,418 |
| HPRD | 38,943 | 9,199 | 3,985 |
| BioGRID | 744,665 | 36,260 | 6,476 |
| STRING | 24,185,620 | 324,767 | 10,320 |
| SIFTS | 27,204 | 23,414 | 6,968 |

For each data source, the PPIs *ppi_k_* were represented by ordered pairs of UniProtIds (*L_k_, R_k_*), in which *L_k_* always precedes *R_k_* in the alphanumerical order to avoid redundancy.

Associations between UniProtIds and Pfam domains were then extracted from UniProtKB/SwissProt and UniProtKB/TrEMBL to give a dataset of UniProtIds-Pfam relationships for instanciating *E*_1_ in 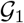 and *E*_2_ in 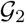 respectively. During this process, it is possible to represent Pfam domains as subsets of size 1 and to generate domain subsets of size 2 in order to infer multiple domain DDIs. However, this extension of PPIDM was not further exploited in this study.

### Methods for PPIDM result evaluation

The set of inferred DDIs was compared with various other sets of DDIs using Python dataframes and a setwise representation for DDIs to avoid duplicate comparison of symmetrical pairs. The June2017, January2018, January2019 and April2020 releases of 3did were used after subtracting the Gold-Standard data set (7, 254 DDIs), in order to check how many inferred DDIs correspond to DDIs in 3did that were not used as Gold-Standard during PPIDM generation, and how many inferred DDIs correspond to DDIs that were not known from 3did at the time PPDIM was generated. The DOMINE 2011 [39] dataset was also downloaded from its web sites (https://manticore.niehs.nih.gov/cgi-bin/Domine and prepared for similar comparison purpose. Venn diagrams were generated using the matplotlib Python library.

The coverage of PPIs by DDIs was measured in a large set of non redundant PPIs, named here IMEx-PPIs, extracted from the curated database assembled by the International Molecular Exchange Consortium (IMEx; [54, 55]), downloaded from https://www.imexconsortium.org/ on 23 August, 2020. Another dataset was also built using the human STRING subset of PPIs [56], downloaded on 30 July, 2019, and filtered to retain only PPIs with a STRING score greater than 800. This dataset is named here hSTRING-PPIs. Only PPIs composed of proteins containing at least one Pfam domain were retained, leading to a total of 75, 865 and 1, 754, 203 PPIs for IMEx-PPIs and hSTRING-PPIs respectively. A PPI is counted as covered by a DDI inferred from PPIDM when at least one DDI from this dataset is present among the pairs of domains derived from the domain composition of each protein in the PPI. A corollary result is the estimation of the percentage of useful DDIs in the dataset. A useful DDI is a DDI used to cover at least one PPI. Counting was performed with PPIDM and DOMINE datasets. A negative control was added here thanks to a dataset of negatively labelled DDIs known as Negatome2.0 [53] that was downloaded from http://mips.helmholtz-muenchen.de/proj/ppi/negatome/), and prepared similarly to the other datasets.

## Results

### Data Source Weights and Similarity Score Threshold

Our merged dataset of scored DDIs contains 513, 260 IntAct, 75, 823 DIP, 97, 487 MINT, 69,940 HPRD, 816,807 BioGRID, 4,131,112 STRING-EXP, 4,050,795 STRING-REST, and 60,114 SIFTS candidate DDIs between Pfam domains, giving a total of 4, 592, 763 distinct DDIs (Table 2). In our ROC-based training procedure, the best AUC value of 0.9944 was obtained with weights *w_IntAct_* = 0.05, *w_DIP_* = 0.01, *w_MINT_* = 0.01, *w_BioGRID_* = 0.09, *w_STRING–Exp_* = 0.12, *w_STRING–Rest_* = 0.06, *w_HPRD_* = 0.17, and *w_SIFTS_* = 1.0. These weights give far greater importance to the candidate interactions in the SIFTS dataset, which is derived from the PDB, compared to those from other databases. This is consistent with the fact that our Gold-Standard positive instances are observed interactions extracted from PDB entries. The optimal score threshold for the consensus similarity matrix was determined according to the *F*1 – *score* calculated on the learning set of positive and negative DDIs described above. A score threshold of 0.01586 was obtained for a maximal F-Measure of 0.9718 on the Training set. Applying this threshold to the Test set yielded a comparable F-measure of 0.9717, and precision and recall values of 0.9808 and 0.9628, respectively.

**Table 2.** Statistics on the source datasets, merged DDIs before filtering and inferred DDIs after filtering at the consensus score threshold.

|  | Name | DDIs | Pfam entries |
| --- | --- | --- | --- |
| Source Datasets | IntAct | 513,260 | 9,898 |
|  | DIP | 75,823 | 6,418 |
|  | MINT | 97,487 | 6,438 |
|  | BioGRID | 816,807 | 6,467 |
|  | STRING-Exp | 4,131,112 | 10,320 |
|  | STRING-Rest | 4,050,795 | 10,313 |
|  | HPRD | 69,940 | 3,985 |
|  | SIFTS | 60,114 | 7,449 |
| Merged DDIs | Before filtering | 4,592,763 | 12,622 |
|  | (Including Gold-Standard) | (7,254) | (5,260) |
| PPIDM | Filtered results | 84,552 | 12,098 |
|  | (Including Gold-Standard) | (6,989) | (5,166) |

### Analysis of the inferred DDIs

The overall results of PPIDM execution are summarized in Table 2. This table shows the numbers of DDIs before and after filtering with the consensus score threshold. After applying the 0.01586 score threshold, the number of pairwise DDIs falls to 84,552 *i.e*. 1.84% of the merged dataset with an overlap of about 96.3% (6, 989 ÷ 7, 254) with the Gold-Standard (3did ⋂ KBDOCK) reference DDIs. Table 3 shows the distribution of PPIDM inferred DDIs in our *Gold, Silver*, and *Bronze* categories, along with the degree of overlap with the Gold-Standard reference dataset. This table shows that PPIDM provides 9, 175 *Gold* DDIs (with only statistically significant scores in more than 4 PPI sources) and 24, 934 *Silver* DDIs (with only statistically significant scores in less than 4 PPI sources), and 50, 443 *Bronze* DDIs (having at least one non statistically significant score). However, it also shows that the 6, 989 Gold-Standard DDIs retrieved in PPIDM are broken down in 2, 852 *Gold*, 3, 888 *Silver* and only 249 *Bronze* DDIs. This is a good indication that the PPIDM scoring and filtering strategy likely infers high quality and relevant new DDIs in the *Gold* and *Silver* categories.

**Table 3.**
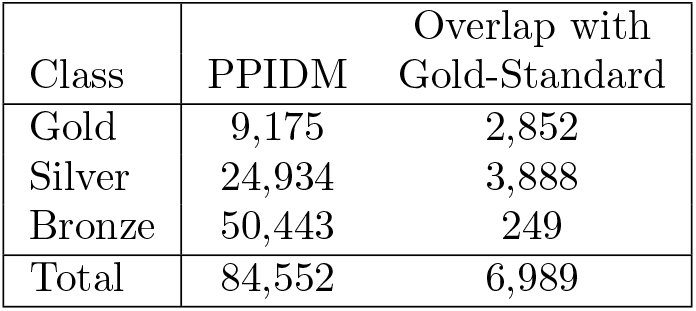
The distribution in *Gold, Silver* and *Bronze* categories of all DDIs inferred by PPIDM, and of their overlap with our Gold-Standard.

### Comparison with the DOMINE dataset

In order to compare PPIDM with the DOMINE dataset, we extracted DDIs from the file available from the latest version of the DOMINE database (2011). The DOMINE file contains 26,219 inferred DDIs (Pfam-Pfam interactions) with 5,410 distinct Pfam domains. This set (shown as green in Fig 2) was compared with the set of all 84, 552 DDIs inferred by PPIDM (blue in Fig 2). This comparison showed that 8, 433 DDIs from DOMINE (about one third of this dataset) are present in the PPIDM dataset, including 4, 824 DDIs from the Gold-Standard (Intersection between yellow, blue, and green in Fig 2).

**Fig 2.**
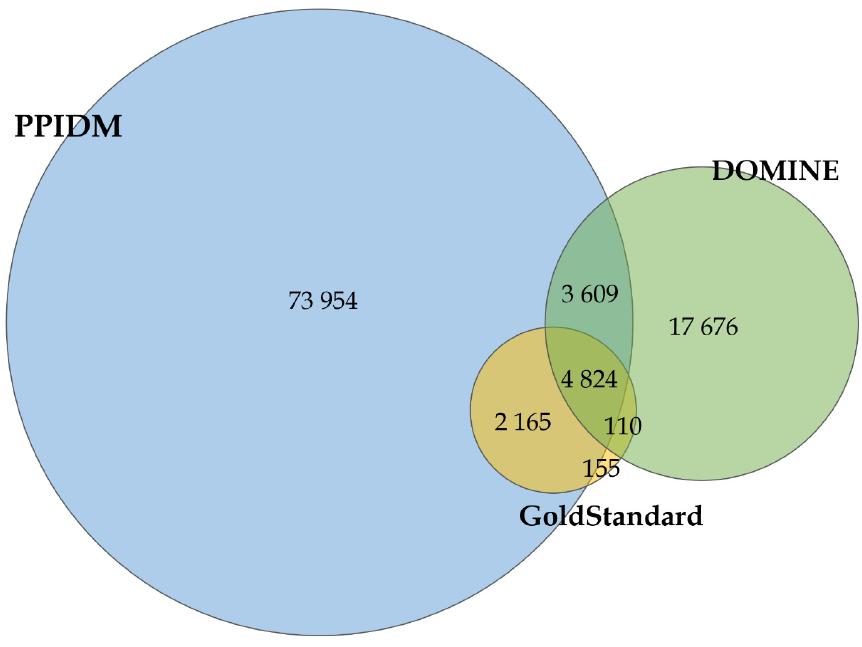
Venn diagram for overlapping DDIs between PPIDM (blue), DOMINE (green), and our Gold-Standard (KBDOCK ⋂ 3did, yellow). PPIDM and DOMINE share 8,433 (3, 609 + 4, 824) DDIs. The Gold-Standard has 6, 989 and 4, 934 DDIs in common with PPIDM and DOMINE, respectively, while the Gold-Standard, PPIDM, and DOMINE share together 4, 824 interactions.

The remaining 17, 676 DDIs in DOMINE were then compared with the DDIs from the Gold-Standard. This comparison (the intersection of purple and yellow minus blue) showed that only 110 DDIs are common to DOMINE and the Gold-Standard but not to PPIDM, indicating that PPIDM misses only 2% (110 ÷ 4,934) of the DOMINE DDIs confirmed by the Gold-Standard. This comparison also shows that the PPIDM result set only infers 3, 609 DOMINE DDIs from its 21,285 (26, 219 – 4, 934) DDIs not shared with the Gold-Standard. This percentage is rather low (around 17%) and likely reflects the heterogeneity of methods used for DDI inference in DOMINE and the lack of consensus between the datasets merged in DOMINE.

### Evaluation of PPIDM Predictions

It is very difficult to review individually the large amount of DDIs inferred by PPIDM. A first attempt of global evaluation has consisted in analyzing the capacity of PPIDM to infer experimental DDIs, for example observed in PDB complexes, outer from the Gold-Standard. For this purpose, we downloaded the DDI sets from the 2017(june), 2018, 2019 and 2020 releases of the 3did database, we subtracted the Gold-Standard set of DDIs (7, 254 DDIs) leading to the so-called 3*did_δ_* dataset. We then checked the overlap between the remaining DDIs and the set of DDIs inferred by PPIDM (Fig 3, panel A) or DOMINE (panel B). It can be seen that the DDIs inferred by PPIDM cover an increasing number of DDIs from the various 3did releases. The percentage of coverage decreases with time (from 70% for the 2017 release to about 46% for the 2020 release). A similar behaviour is observed with the DOMINE dataset but to a much lesser extend. The percentage of coverage varies from 28% for the 2017 release, to about 17% for the 2020 release of 3did, at the advantage of the PPIDM approach. More precisely, from the 3, 854 new DDIs in 3did 2020 that were unknown in 2017 (when substracting 3did2017(february) from 3did2020), PPIDM has inferred exactly 800 DDIs and DOMINE only 234.

**Fig 3.**
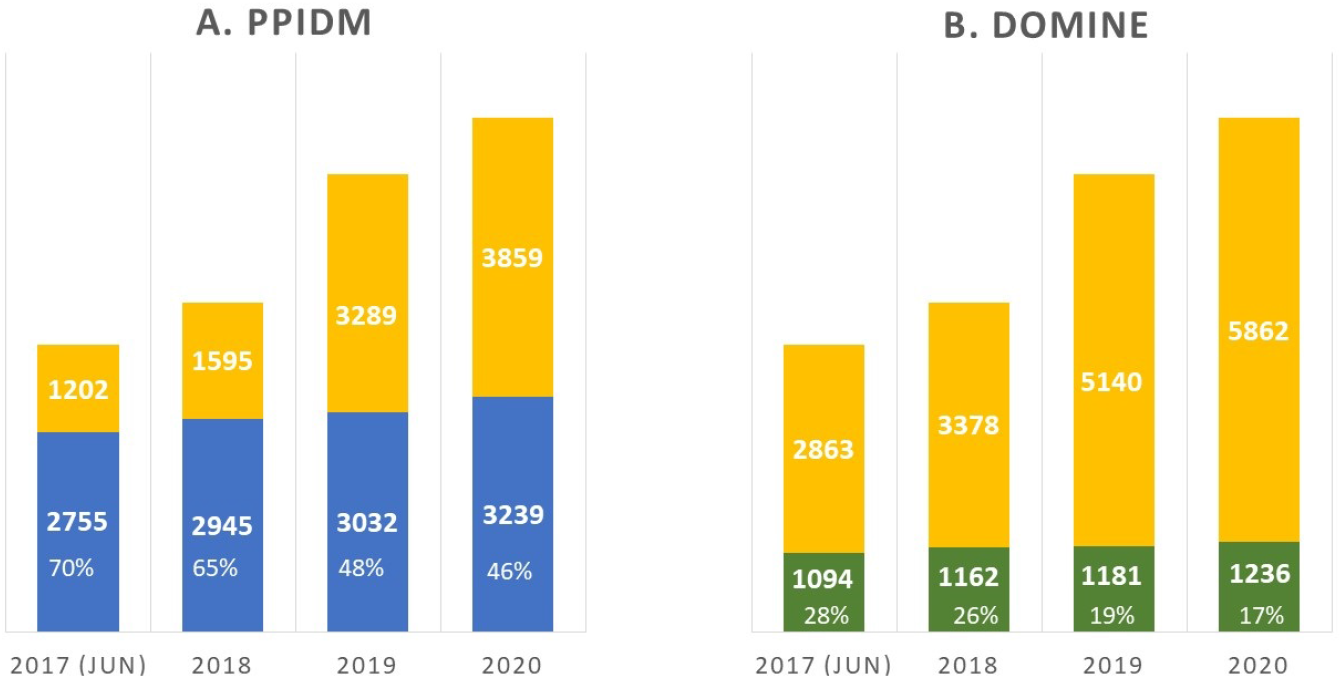
Coverage of 3*did_δ_* by inferred DDIs from PPIDM (blue; panel A) and DOMINE (green; panel B).

The disadvantage of DOMINE here is likely due to the fact that the datasets it contains have been produced several years ago (from 2004 to 2010). However for PPIDM, it shows that a non negligeable subset of inferred DDIs are validated by observations in 3did that did not exist at the time PPIDM was produced.

The decrease in the percentage of coverage of 3did by PPIDM inferred DDIs can be explained by the rapid growth of 3did that directly follows the growth of the PDB. Recent PDB entries, posterior to 2017 could not be used by our version of PPIDM (2017). Indeed, we determined that among the 3, 859 DDIs not inferred by PPIDM in 3did 2020, 90.3% had a very limited number (*Nb* ≤ 5) of PDB entries before 2017, including 39.1% of DDIs without any PDB entry in 2017 or before. By design, it was impossible for PPIDM to infer such DDIs.

Another way to evaluate the PPIDM set of inferred DDIs was to study the coverage of existing PPIs by these DDIs. For this purpose, we used a recent dataset (august 2020) of 50,032 non redundant curated PPIs from the IMEX database [55]. Only PPIs composed of proteins containing at least one Pfam domain were retained to allow coverage analysis by 3did 2020, PPIDM, DOMINE, and Negatome as a negative control. Results are presented in Fig 4A and reveal that PPIDM covers 13, 316 PPIs from IMEx, *i.e*. about 4,000 PPIs more than DOMINE and almost 2.2 times higher than the number covered by 3did 2020.

**Fig 4.**
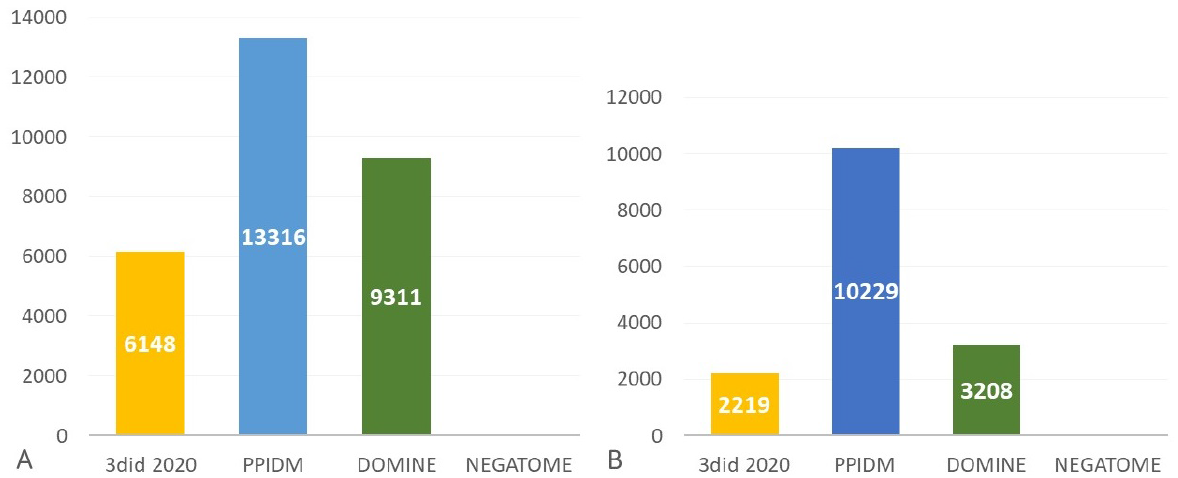
Coverage of PPI interactome derived from IMEx database (50, 032 PPIs). (A) Number of PPIs covered by at least one DDI; (B) Number of DDIs covering at least one PPI.

To complement these results, we also computed the number of DDIs that cover at least one PPI of the IMEx dataset. The obtained results are shown in Fig 4B. The number of relevant DDIs for PPI coverage in PPIDM is more than 3 times the number of relevant DDIs in DOMINE and 4.6 times higher than the number of relevant DDIs in 3did2020. Thus, the PPIDM dataset encompasses an interesting diversity of DDIs that deserves further studies.

Similar results were obtained at a larger scale using a subset of STRING containing 607, 088 human well-scored PPIs (Fig 5). These results yield an update of the capacity of existing available sets of DDIs to cover large-scale sets of PPIs on the basis of their Pfam composition : about 24%, 35% and 30% of the STRING subset of PPIs are covered by DDIs from 3did2020, PPIDM and DOMINE, respectively (Fig 5A). Moreover, a total of 23,297 DDIs (*i.e*. 27.5% of the PPIDM dataset) are used to cover at least one PPI of the hSTRING-PPIs dataset (Fig 5B).

**Fig 5.**
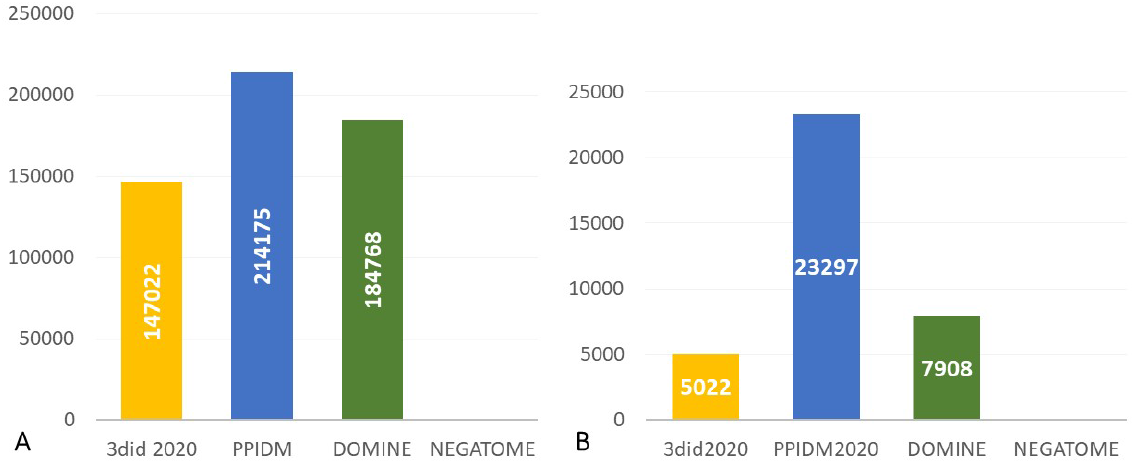
Coverage of human interactome derived from STRING (607, 088 PPIs). (A) Number of PPIs covered by at least one DDI; (B) Number of DDIs covering at least one PPI.

Finally we selected a few examples of newly inferred DDIs to study their possible biological interest. We selected the PPIDM (Gold) DDIs that cover at least one PPI from Imex and filtered out the DDIs present in 3did 2020 or DOMINE. There were 1,915 DDIs left. We extracted those DDIs covering more than 10 PPIs in IMEx (155 DDIs) and ranked them from higher to lower PPIDM consensus score. in Table 4, we present five examples selected on the basis of interpretable Pfam description among the top-15 DDIs.

**Table 4.** Description of selected *Gold* DDIs newly inferred by PPIDM and absent from 3did2020 or DOMINE. The PPIDM consensus score and the number of covered PPIs in the IMEx database are indicated. The domain short name is taken from the corresponding Pfam entry and followed by a short description adapted from the corresponding Pfam entry.

| Inferred DDI | PPIDM score | Covered PPIs | Domain 1 | Domain 2 |
| --- | --- | --- | --- | --- |
| PF01352-PF01352 | 0.456 | 31 | KRAB box: the KRAB domain (or Kruppel-associated box) is present in about a third of zinc finger proteins containing C2H2 fingers. The KRAB domain is found to be involved in protein-protein interactions. | KRAB box: see left. |
| PF11629-PF16517 | 0.315 | 11 | Mst1_SARAH: this family of proteins represents the C terminal SARAH domain of Mst1. | Nore1-SARAH: Nore1 forms homo- and hetero complexes through its C-terminal SARAH domain. |
| PF03931-PF12937 | 0.252 | 26 | Skp1_POZ: Skp1 family, tetramerisation domain. This domain is found at the N-terminal of SKP1 proteins. | F-box-like: Through the F-box, proteins are linked to the SKP1 protein. |
| PF00622-PF13765 | 0.202 | 15 | SPRY: SPRY Domain is named from SPLa and the RYanodine receptor. It is a protein-interaction module involved in many important signalling pathways. | PRY: a 50-60 amino acids domain associated with SPRY domains, adjacent to its N-terminal. PRY and SPRY domains are structurally very similar and consist of a beta sandwich fold. |
| PF00104-PF11825 | 0.184 | 12 | Hormone_recep: this all helical domain is involved in binding the hormone in these receptors. | Nuc_recep-AF1: Nuclear/hormone receptor activator site AF-1. |

Only the first DDI here is an homodimer. The concerned domain PF01352 is part of a zinc finger protein and is involved in protein-protein interactions. The observation of this inferred DDI suggests that at least some zinc finger proteins could act as a dimer. The second DDI in the table (PF11620-PF16517) involves two domains sharing the same SARAH region which is reported to mediate the formation of homo and hetero complexes. The third DDI (PF03931-PF12937) is beween a domain present in SKP1 proteins and a domain (F-Box-like) known to mediate interaction with SKP1 proteins. Our fourth example (PF00622-PF13765) appears to concern two frequently associated domains PRY and SPRY, that are very similar. The SPRY domain is reported as a protein-interaction module but it remains to be demonstrated whether the two domains actually interact together. Finally, the last DDI in the table (PF00104-PF11825) involves the hormone-binding and activator-site domains of nuclear receptors. It is well known that these two domains are close to each other on nuclear receptors and they are likely to interact because binding to hormones often induces allosteric conformational changes leading to binding of an activator.

More examples of relevant new DDIs can be found on the website displaying all *Gold* DDIs as a searchable table by Pfam identifier (http://ppidm.loria.fr).

## Discussion

This paper has introduced and presented a new approach named PPIDM for mining protein-protein interactions at the domain level. Threshold optimization was carried out with a Gold-Standard of positive and negative examples and yielded very high F-measure corresponding to high recall and precision values. The resulting dataset contains 84, 552 inferred DDIs overlapping with 96.3% of the Gold-Standard positive examples derived from 3did. The PPIDM dataset of inferred DDIs represents the largest available repository of inferred DDIs to date, that also includes the largest number of DDIs involved in explaining a large number of non redundant curated PPIs.

The large size of the PPIDM dataset contrasts with the more limited sizes of DOMINE or DIMA 3.0 datasets (around 26 and 50 thousands of DDIs respectively). However, other authors already got even larger datasets in the past. In [9], a dataset of 288, 098 DDIs was inferred using two neighborhood cohesiveness coefficients and a combined proportion method on the STRING interactome. In that work as in ours, the purpose is not the same as with the parsimonious methods that aim to explain a maximum number of PPIs in an interactome with the minimum number of DDIs. Indeed, large repositories of DDIs should rather be understood as reservoirs from which possible DDIs can be picked up to support working hypotheses or to select candidate interactants for a protein of interest.

As the DDIs inferred by PPIDM are qualified with a score and a p-value, many explorations are possible from now on with this dataset. For example it will be interesting to analyse the 800 DDIs inferred in 2017 that were subsequently found new in 3did between 2017 and 2020, in order to learn which features or annotations associated with these DDIs have favored their experimental confirmation in 3did.

Network properties have been computed using the Cytoscape software for PPIDM Gold and Silver subsets of DDI and are compared with the results obtained for 3did 2020 and DOMINE. Comparison is also performed with the two PPI networks used for PPIDM evaluation. Table 5 summarizes the comparison in terms of connected components and node degree distribution. All the networks considered here display the global organisation already reported in [40]: a giant component without any detectable subcluster and a series of smaller isolated components. However the percentage of nodes participating in the giant component is much higher in the PPI networks (more than 97 %) than in the DDI networks (less than 75 %). Consequently, DDI networks contain many more small connected components than PPI networks, including numerous components of size 1 corresponding to self-interacting domains, and components of size 2 corresponding to isolated DDIs. This difference in the distribution of connected components is consistent with the notion that PPIs can display much more interconnections than DDIs because proteins can interact with multiple partners via several domains.

**Table 5.** Comparison of network statistics for (A) DDI networks: 3did2020, PPIDM (Gold), PPIDM (Silver), DOMINE, and (B) PPI networks used for PPIDM evaluation: IMEX and hSTRING. * Degree Sup is the superior degree value for 99% of the nodes ranked according to increasing degree. ** PfamId or UniProtId corresponding to the nodes having the maximal degree. PF07686: Immunoglobulin V-set domain; PF00018: SH3 domain; PF00096: Zinc finger, C2H2 type, domain; PF00069: PKinase domain; A8MQ03: Cysteine-rich tail protein 1; Q09472: Histone acetyltransferase p300.

|  | A. DDIs |  |  |  | B. PPIs |  |
| --- | --- | --- | --- | --- | --- | --- |
|  | 3did 2020 | PPIDM (Gold) | PPIDM (Silver) | DOMINE | IMEx | hSTRING |
| Nodes | 8,048 | 3,771 | 9,195 | 5,410 | 10,016 | 15,472 |
| Edges | 14,278 | 9,175 | 24,934 | 26,219 | 50,031 | 607,087 |
| Connected components (CC) analysis |  |  |  |  |  |  |
| Total CCs | 2,879 | 1,123 | 1,750 | 1,524 | 140 | 40 |
| Giant CC size (nodes) | 3,936 | 1,822 | 6,880 | 3,877 | 9,779 | 15,407 |
| % nodes in giant CC | 48.9 % | 48.3 % | 74.8 % | 71.7 % | 97.6 % | 99.6 % |
| 1-node CCs | 2,244 | 727 | 1,368 | 1,052 | 57 | 23 |
| 2-node CCs | 404 | 239 | 271 | 143 | 69 | 12 |
| Degree distribution (number of nodes) analysis |  |  |  |  |  |  |
| Degree 1 | 1,110 | 445 | 1,248 | 722 | 2,496 | 1,189 |
| Degree 2 | 2,807 | 894 | 2,154 | 1,401 | 1,466 | 917 |
| Degree 3 | 1,683 | 720 | 1,419 | 708 | 1,047 | 702 |
| % nodes with degree > 3 | 30.4 % | 45.4 % | 47.6 % | 47.7 % | 50.0 % | 81.9 % |
| Degree Sup* | 17 | 27 | 30 | 98 | 78 | 431 |
| Degree Max | 368 | 88 | 108 | 499 | 451 | 2,423 |
| PfamId or UniProtId** | PF07686 | PF00018 | PF00096 | PF00069 | A8MQ03 | Q09472 |

It should be noted that the number of connected components of size 1 correspond to isolated domains that only interact with themselves. However self-interacting domains (or homo-DDIs) are also present in the other components. In total PPIDM counts 7,203 homo-DDIs of which exactly 3000 and 4160 are present in the Gold and Silver subsets respectively, corresponding to 79.6% and 45.2 % of the nodes respectively. These percentages can be compared to those observed in 3did 2020 (70.6 %), DOMINE (64.2 %), or the one reported for UniDomInt (32.5 %) in [40]. This clearly indicates that not all domains are capable of self interactions and that, in terms of network properties, the PPIDM (Gold) dataset is closer to the reference 3did 2020 than are the PPIDM (Silver) or DOMINE datasets.

The distribution of nodes according to their degree is also different between DDI and PPI networks. The general shape is right-skewed with a large number of nodes having small degrees and unique nodes with very high degrees (hubs). This confirms that none of these networks is random but, on the contrary, they represent complex sets of interactions. In the four DDI networks considered in this study, the maximum number of nodes is always obtained for degree 2 whereas in the two PPI networks it is for degree 1. The superior degree for 99% of the ranked nodes is rather low for DDI networks (17, 27 and 30 for 3did 2020, PPIDM (Gold) and PPIDM (Silver) respectively), except for DOMINE (98). This superior degree is very high for hSTRING (431) and intermediate for IMEX (78). The nodes with the maximal degree are also indicated in Table 5 and reach very high values for PPI networks. In DDI networks, these outliers indeed correspond to domains responsible for interactions in a large variety of proteins. Three of them (PF00018, PF00069 and PF00096) belong to the list of outliers observed in UniDomInt [40].

A limit of this study is certainly that the PPIDM dataset has been produced on 2017 releases of available PPI resources. While it is always possible to update the dataset on more recent releases, it seemed relevant to us to explore the predictive value of these results derived from previous sources in the light of new data (PPIs or DDIs) obtained subsequently.

Another limit lies in the fact that the PPI sources used in this study are overlapping. For example, STRING imports its experimental data from IMEx, BioGRID and PDB, among other sources and IMEx is made up of IntAct, MINT and DIP. Thus, a non negligeable proportion of PPIs have been reused several times to infer DDIs. Nevertheless, it should be noted that the CODAC process computes similarity scores and p-values for each PPI source separately. These separated values are available in the datasets proposed for download at http://ppidm.loria.fr. Then, data reuse is only effective (and somewhat inevitable) when merging scores, as in the case of other combined DDI datasets such as DOMINE. Another study would be to estimate the impact of this reuse of PPIs on the number or quality of DDIs recovered. Reuse of PPIs also occurs at the evaluation step with IMEx and a subset of STRING. However, this evaluation step does not claim to be similar to the validation of a predictive model in machine learning. It is more akin to evaluating the quality of a corpus produced by an information retrieval procedure. Indeed, the main outcome of our work is to produce a large reservoir of DDIs available for further studies.

Just like all other studies dealing with DDIs, PPIDM encounters by design a limitation in explaining PPIs that do not interact through well-defined conserved domains but rather through intrinsically disordered regions [58].This is likely the reason why a certain percentage of the interactome cannot be explained by inferred DDIs even with a large dataset such as PPIDM.

With the tripartite graph paradigm, our PPIDM inference method is related to link prediction methods in multipartite or knowledge graphs. Our generic graph formulation of link prediction has been found to be useful for modeling the multi-source approach. In addition, the PPIDM setting makes it possible to infer interactions with domains subsets in addition to single domain interactions. This is not the case for most other DDI-inferring methods, except for the discriminative feature selection method described in [30]. For PPIDM, we just need to consider subsets of domain as elements of the two sets *X* and *Y* in the tripartite framework and define their relation to PPIs in *Z*. This extension of PPIDM is quite original and could provide clues in the analysis of multi-domain protein interactions. Indeed, while a small number of single domain proteins interact with their biological associates directly, a much larger number of proteins have more than one domain [59], and interactions between these multi-domain proteins can often involve two or more domains [60].

## Conclusion

This paper has introduced and presented a new approach named PPIDM for inferring DDIs from multiple sources of PPIs. The obtained dataset displays interesting properties including a good scoring of experimental DDIs and a significant coverage of existing PPIs. Further studies will be devoted to better describe the inferred DDIs and possibly validate them through large-scale experimental studies. It could be envisaged for instance to look for correspondences between PPIDM DDIs and cross-linked peptides obtained with cross-linked mass spectrometry, as already performed for PPIs in *Drosophila* embryos [61]. We believe that the large set of inferred DDIs produced in this study can be used to interpret interactome data and provide added-value to protein network analyses.

## Acknowledgements

This article is dedicated to David W. Ritchie who passed away in September 2019, in memory of his interest in this work and in tribute to his many scientific contributions in structural bioinformatics. This work and the publication of this article were funded by Agence Nationale de la Recherche (grant numbers: ANR-11-MONU-006-02 and ANR-15-RHUS-0004), Inria Nancy Grand Est, and FEDER-Region Grand-Est (CPER IT2MP). SZA was recipient of an Inria CORDI-S PhD fellowship, AAN benefited from a post-doctoral fellowship funded by the Region Grand-Est and the Faculty Hospital of Nancy, and MDD from an interface contract between CNRS and the Faculty Hospital of Nancy.

## Data availability statement

All code written in support of this publication is publicly available at https://gitlab.inria.fr/capsid.public_codes/ppidmpublic.

Input files and generated data are available from https://doi.org/10.5281/zenodo.4880347.

## References

1. El-Gebali S, Mistry J, Bateman A, Eddy SR, Luciani A, Potter SC, et al. The Pfam protein families database in 2019. Nucleic Acids Research. 2019;47(Database Issue):D427–D432. doi:10.1093/nar/gky995.

2. Marchler-Bauer A, Yu B, Han L, He J, Lanczycki CJ, Lu S, et al. CDD/SPARCLE: functional classification of proteins via subfamily domain architectures. Nucleic Acids Research. 2017;45(Database Issue):D200–D203. doi:10.1093/nar/gkw1129.

3. Stein A, Russell RB, Aloy P. 3did: interacting protein domains of known three-dimensional structure. Nucleic Acids Research. 2005;33(Database Issue):D413–D417.

4. Stein A, Céol A, Aloy P. 3did: identification and classification of domain-based interactions of known three-dimensional structure. Nucleic Acids Research. 2011;39(Database Issue):718–723. doi:10.1093/nar/gkq962.

5. Mosca R, Céol A, Stein A, Olivella R, Aloy P. 3did: a catalog of domain-based interactions of known three-dimensional structure. Nucleic Acids Research. 2014;42(Database Issue):374–379. doi:10.1093/nar/gkt887.

6. Ghoorah AW, Devignes MD, Smaïl-Tabbone M, Ritchie DW. KBDOCK 2013: a spatial classification of 3D protein domain family interactions. Nucleic Acids Research. 2013;42(Database Issue):D389–D395.

7. Finn RD, Miller BL, Clements J, Bateman A. iPfam: a database of protein family and domain interactions found in the Protein Data Bank. Nucleic Acids Research. 2013;42(Database Issue):D364–D373.

8. Meyer MJ, Das J, Wang X, Yu H. INstruct: a database of highquality 3D structurally resolved protein interactome networks. Bioinformatics. 2013;29(12):1577–1579.

9. Segura J, Sorzano COS, Cuenca-Alba J, Aloy P, Carazo JM. Using neighborhood cohesiveness to infer interactions between protein domains. Bioinformatics. 2015;31(15):2545–2552.

10. Sprinzak E, Margalit H. Correlated sequence-signatures as markers of protein-protein interaction. Journal of molecular biology. 2001;311(4):681–692.

11. Kim W, Park J, Suh J. Large scale statistical prediction of protein-protein interaction by potentially interacting domain (PID) pair. Genome Informatics. 2002;13:42–50.

12. Nye TM, Berzuini C, Gilks WR, Babu MM, Teichmann SA. Statistical analysis of domains in interacting protein pairs. Bioinformatics. 2004;21(7):993–1001.

13. Deng M, Mehta S, Sun F, Chen T. Inferring domain–domain interactions from protein–protein interactions. Genome research. 2002;12(10):1540–1548.

14. Riley R, Lee C, Sabatti C, Eisenberg D. Inferring protein domain interactions from databases of interacting proteins. Genome biology. 2005;6(10):R89.

15. Wang H, Segal E, Ben-Hur A, Li Q, Vidal M, Koller D. InSite: a computational method for identifying protein-protein interaction binding sites on a proteome-wide scale. Genome Biology. 2007;8(9):R192.

16. Rhodes DR, Tomlins S, Varambally S, Mahavisno V, Barrette T, Kalyana-Sundaram S, et al. Probabilistic model of the human protein-protein interaction network. Nat Biotechnol. 2005;23(8):951–959.

17. Lee H, Deng M, Sun F, Chen T. An integrated approach to the prediction of domain-domain interactions. BMC bioinformatics. 2006;7(1):269.

18. Liu M, Chen X, Jothi R. Knowledge-guided inference of domain-domain interactions from incomplete protein-protein interaction networks. Bioinformatics. 2009;25(19):2492–2499.

19. Ng S, Zhang Z, Tan S, Lin K. InterDom: a database of putative interacting protein domains for validating predicted protein interactions and complexes. Nucleic Acids Research. 2003;31(1):251–254.

20. Pagel P, Wong P, Frishman D. A domain interaction map based on phylogenetic profiling. Journal of molecular biology. 2004;344(5):1331–1346.

21. Jothi R, Cherukuri PF, Tasneem A, Przytycka TM. Co-evolutionary analysis of domains in interacting proteins reveals insights into domain–domain interactions mediating protein–protein interactions. Journal of molecular biology. 2006;362(4):861–875.

22. Pazos F, Valencia A. Protein co-evolution, co-adaptation and interactions. EMBO J. 2008;27(20):2648–2655.

23. Luo Q, Pagel P, Vilne B, Frishman D. DIMA 3.0: Domain Interaction Map. Nucleic Acids Research. 2011;39(Database Issue):D724–D729.

24. Croce G, Gueudre T, Ruiz-Cuevas M, Keidel V, Figliuzzi M, Szurmant H, et al. A multi-scale coevolutionary approach to predict interactions between protein domains. PLoS Comput Biol. 2019;15(10):e1006891.

25. Guimarães KS, Jothi R, Zotenko E, Przytycka TM. Predicting domain-domain interactions using a parsimony approach. Genome biology. 2006;7(11):R104.

26. Guimarães KS, Przytycka TM. Interrogating domain-domain interactions with parsimony based approaches. BMC Bioinformatics. 2008;9:171.

27. Chen C, Zhao JF, Huang Q, Wang RS, Zhang XS. Inferring domain-domain interactions from protein-protein interactions in the complex network conformation. BMC systems biology. 2012;6(1):S7.

28. Singhal M, Resat H. A domain-based approach to predict protein-protein interactions. BMC Bioinformatics. 2007;8:199.

29. Memisevic V, Wallqvist A, Reifman J. Reconstituting protein interaction networks using parameter-dependent domain-domain interactions. BMC Bioinformatics. 2013;14:154.

30. Chen XW, Liu M. Prediction of protein–protein interactions using random decision forest framework. Bioinformatics. 2005;21(24):4394–4400.

31. Zhao XM, Chen L. Domain-Domain Interaction Identification with a Feature Selection Approach. Lecture Notes in Bioinformatics. 2008;5265:178–186.

32. Zhao X, Chen L, Aihara K. A discriminative approach for identifying domain-domain interactions from protein-protein interactions. Proteins. 2010;78(5):1243–1253.

33. Khor S. Inferring domain-domain interactions from protein-protein interactions with formal concept analysis. PloS one. 2014;9(2):e88943.

34. Salwinski L, Miller CS, Smith AJ, Pettit FK, Bowie JU, Eisenberg D. The Database of Interacting Proteins: 2004 update. Nucleic Acids Research. 2004;32(Database Issue):D449–D451. doi:https://doi.org/10.1093/nar/gkh086.

35. Kerrien S, Aranda B, Breuza L, Bridge A, Broackes-Carter F, Chen C, et al. The IntAct molecular interaction database in 2012. Nucleic Acids Research. 2012;40(Database Issue):D841–D846. doi:https://doi.org/10.1093/nar/gkr1088.

36. Keshava Prasad TS, Goel R, Kandasamy K, Keerthikumar S, Kumar S, Mathivanan S, et al. Human Protein Reference Database–2009 update. Nucleic Acids Research. 2009;37(Database Issue):D767–D772. doi:https://doi.org/10.1093/nar/gkn892.

37. Szklarczyk D, Morris JH, Cook H, Kuhn M, Wyder S, Simonovic M, et al. The STRING database in 2017: quality-controlled protein-protein association networks, made broadly accessible. Nucleic Acids Research. 2017;45(Database Issue):D362–D368. doi:https://doi.org/10.1093/nar/gkw937.

38. Raghavachari B, Tasneem A, Przytycka TM, Jothi R. DOMINE: a database of protein domain interactions. Nucleic Acids Research. 2007;36(Database Issue):D656–D661.

39. Yellaboina S, Tasneem A, Zaykin DV, Raghavachari B, Jothi R. DOMINE: a comprehensive collection of known and predicted domain-domain interactions. Nucleic Acids Research. 2010;39(Database Issue):D730–D735.

40. Björkholm P, Sonnhammer EL. Comparative analysis and unification of domain-domain interaction networks. Bioinformatics. 2009;25(22):3020–3025.

41. Kim Y, Min B, Yi G. IDDI: The Integrated Domain-Domain Interaction Analysis System. In: Wu F, Zaki MJ, Morishita S, Pan Y, Wong S, Christianson A, et al., editors. IEEE International Conference on Bioinformatics and Biomedicine, BIBM 2011, Atlanta, GA, USA, November 12-15,, 2011. IEEE Computer Society; 2011. p. 520–525.

42. Alborzi SZ, Ritchie DW, Devignes M. Computational discovery of direct associations between GO terms and protein domains. BMC Bioinformatics. 2018;19-S(14):53–66.

43. Cui X, Churchill GA, et al. Statistical tests for differential expression in cDNA microarray experiments. Genome Biol. 2003;4(4):210.

44. Mogotsi I. Manning, Christopher D. and Raghavan, Prabhakar and Schütze, Heinrich : Introduction to information retrieval; 2010.

45. Fawcett T. An introduction to ROC analysis. Pattern recognition letters. 2006;27(8):861–874.

46. Chawla NV, Japkowicz N, Kotcz A. Special issue on learning from imbalanced data sets. ACM Sigkdd Explorations Newsletter. 2004;6(1):1–6.

47. Orchard S, Ammari M, Aranda B, Breuza L, Briganti L, Broackes-Carter F, et al. The MIntAct project–IntAct as a common curation platform for 11 molecular interaction databases. Nucleic Acids Research. 2014;42(Database Issue):D358–D363. doi:https://doi.org/10.1093/nar/gkt1115.

48. Chatr-Aryamontri A, Oughtred R, Boucher L, Rust J, Chang C, Kolas NK, et al. The BioGRID interaction database: 2017 update. Nucleic Acids Research. 2016;45(Database Issue):D369–D379. doi:https://doi.org/10.1093/nar/gkw1102.

49. Oughtred R, Rust J, Chang C, Breitkreutz BJ, Stark C, Willems A, et al. The BioGRID database: A comprehensive biomedical resource of curated protein, genetic, and chemical interactions. Protein Sci. 2021;30(1):187–200.

50. Velankar S, Dana JM, Jacobsen J, van Ginkel G, Gane PJ, Luo J, et al. SIFTS: Structure Integration with Function, Taxonomy and Sequences resource. Nucleic Acids Research. 2013;41(Database Issue):D483–D489. doi:https://doi.org/10.1093/nar/gks1258.

51. Dana JM, Gutmanas A, Tyagi N, Qi G, O’Donovan C, Martin M, et al. SIFTS: updated Structure Integration with Function, Taxonomy and Sequences resource allows 40-fold increase in coverage of structure-based annotations for proteins. Nucleic Acids Res. 2019;47(D1):D482–D489.

52. Ghoorah AW, Devignes MD, Smaïl-Tabbone M, Ritchie DW. Spatial clustering of protein binding sites for template based protein docking. Bioinformatics. 2011;27(20):2820–2827.

53. Blohm P, Frishman G, Smialowski P, Goebels F, Wachinger B, Ruepp A, et al. Negatome 2.0: a database of non-interacting proteins derived by literature mining, manual annotation and protein structure analysis. Nucleic Acids Research. 2014;42(Database Issue):D396–D400.

54. Orchard S, Kerrien S, Abbani S, Aranda B, Bhate J, Bidwell S, et al. Protein interaction data curation: the International Molecular Exchange (IMEx) consortium. Nature methods. 2012;9(4):345–350. doi:https://doi.org/10.1038/nmeth.1931.

55. Porras P, Barrera E, Bridge A, Del-Toro N, Cesareni G, Duesbury M, et al. Towards a unified open access dataset of molecular interactions. Nat Commun. 2020;11(1):6144.

56. Szklarczyk D, Gable AL, Lyon D, Junge A, Wyder S, Huerta-Cepas J, et al. STRING v11: protein-protein association networks with increased coverage, supporting functional discovery in genome-wide experimental datasets. Nucleic Acids Research. 2019;47(Database Issue):D607–D613. doi:https://doi.org/10.1093/nar/gky1131.

57. Allen MD, Bycroft M, Zinzalla G. Structure of the BRK domain of the SWI/SNF chromatin remodeling complex subunit BRG1 reveals a potential role in protein–protein interactions. Protein Science. 2020;29(4):1033–1039. doi:https://doi.org/10.1002/pro.3820.

58. Fong JH, Shoemaker BA, Garbuzynskiy SO, Lobanov MY, Galzitskaya OV, Panchenko AR. Intrinsic Disorder in Protein Interactions: Insights From a Comprehensive Structural Analysis. PLoS Comput Biol. 2009;5(3). doi:10.1371/journal.pcbi.1000316.

59. Apic G, Gough J, Teichmann SA. Domain combinations in archaeal, eubacterial and eukaryotic proteomes. Journal of molecular biology. 2001;310(2):311–325.

60. Bhaskara RM, Srinivasan N. Stability of domain structures in multi-domain proteins. Scientific reports. 2011;1.

61. Götze M, Iacobucci C, Ihling CH, Sinz A. A Simple Cross-Linking/Mass Spectrometry Workflow for Studying System-wide Protein Interactions. Analytical Chemistry. 2019;91(15):10236–10244. doi:10.1021/acs.analchem.9b02372.

